# Extended poly(A) tails are a shared feature of herpesvirus mRNAs

**DOI:** 10.64898/2025.12.15.694445

**Authors:** Erik Fuhrmann, Sae Toda, Jonas Leins, Pierina Cetraro, Vedang Deshpande, Carina Jacobsen, Kai A. Kropp, Mart M. Lamers, Elene Loliashvili, Mostafa Saleban, Ruth Verstraten, Carolin Vogt, Wiyada Wongwiwat, Werner J.D. Ouwendijk, Abel Viejo-Borbolla, Robert E. White, Angus C. Wilson, Hannah M. Burgess, Daniel P. Depledge

## Abstract

Poly(A) tails are present on most cellular and viral mRNAs, providing a platform for poly(A)-binding proteins that stimulate translation and regulate the deadenylation and stability of transcripts in the cytoplasm. Here we leverage nanopore direct RNA sequencing to analyse the distribution of poly(A) tail lengths on cellular and viral mRNAs across Herpesviridae and other DNA and RNA virus infections. We find that herpesvirus mRNA poly(A) tails are consistently longer than those on cellular and other viral transcripts, presenting a previously unrecognized yet widespread mechanism to advantage herpesviral gene expression. This contrasts with the templated poly(A) tails on coronavirus RNAs and those on cytoplasmically transcribed poxviral mRNAs, which are more similar in length to those on host mRNAs. Herpesviral noncoding RNAs display differential poly(A) tailing patterns which do not correlate with nuclear localisation while individual herpesviral mRNAs also show variation in the extent to which their poly(A) tail lengths change during the virus lifecycle, suggestive of additional uncharacterised layers of poly(A) tail length regulation. Importantly, while we detect non-adenosine nucleotides within herpesviral poly(A) tails, which are known to oppose deadenylase activity, this “mixed tailing” is not at sufficient frequency to explain the widespread extended tails of herpesvirus mRNAs.

## Introduction

All viruses are dependent on the host cell translation machinery for viral protein production and have thus evolved strategies to polyadenylate viral RNAs. These include (i) utilization of cellular transcription machinery in the case of nuclear-replicating DNA viruses (e.g. herpesviruses, adenoviruses, and polyomaviruses (1)), (ii) producing a viral poly(A) polymerase (e.g. poxviruses (2)), and (iii) templated transcription of poly(A) tails from an antisense uridine tract (e.g. coronaviruses (3, 4), picornaviruses (5), caliciviruses (6)).

The addition of poly(A) tails to cellular mRNAs and select non-coding RNAs (ncRNAs)during 3’ end processing is important for their stability, export, translation, and decay (1, 7). In mammalian cells, poly(A) polymerase (PAP) adds a ∼200-nt poly(A) tail after transcription (8), with this length dictated by the nuclear poly(A)-binding protein, PABPN1 (9). Over time, the poly(A) tail is trimmed by cytosolic deadenylase complexes PAN2-PAN3 and CCR4-NOT, with full deadenylation believed to commit an mRNA to degradation by decapping and exonucleolytic decay (1). Thus, the bulk population of cellular mRNAs typically possesses mRNA poly(A) tail lengths of 30-100 nt (10–12). The effects of poly(A) tail length on mRNA stability and translation are mediated by cytoplasmic poly(A)-binding proteins (PABPC) of which PABPC1 is the most abundant in mammalian cells. A longer poly(A) tail provides a greater platform for PABPCs, which are able to bind cooperatively (13) and have a ∼27 nucleotide footprint each (14). PABPC at the 3’ poly(A) tail interacts with eIF4G bound at the 5’ cap to promote translation initiation (15). Paradoxically, PABPC1 protects the poly(A) tail from deadenylation, but also helps recruit deadenylases to effect mRNA decay (1). Recruitment of the deadenylase CCR4-NOT by adaptor RNA-binding proteins also brings about targeted degradation of specific mRNAs. Conversely, readenylation by cytoplasmic PAPs are thought to enhance mRNA stability and allow further rounds of translation (16). The abundance of PABPC1 in specific cell types however is speculated to dictate how responsive translation and stability of mRNAs are to poly(A) tail length (17). Importantly, poly(A) tailed viral RNAs will also be subject to regulation through these processes.

While northern blotting (18) and modified short-read sequencing approaches such as PAL-seq and TAIL-seq (10, 19) are favoured methods for measuring poly(A) tail lengths, these have now been complemented by nanopore direct RNA sequencing (DRS, (20)), a methodology which eschews reverse transcription and amplification steps that can introduce biases. DRS sequences native RNAs by passing them through biological nanopores in a 3’-5’ direction and converts associated changes in ionic current into sequence data, allowing estimates of poly(A) tail lengths to be generated using tools such as *nanopolish* (21), *tailfindr* (22), and *boostnano* (23). Crucially, these tools produce near-identical results at very high accuracy when applied to *in vitro* transcribed RNA standards with defined poly(A) tail lengths (23). Moreover, another recently developed tool, *ninetails* interprets ionic current changes within poly(A) tails to resolve the individual positions and identities of non-adenosine nucleotide incorporations (24), a process known as mixed tailing.

Utilizing nanopore DRS we recently determined that poly(A) tails on HCMV RNAs are significantly longer than those on cellular RNAs and are relatively insensitive to deadenylation via CCR4-NOT (25). Intriguingly, HCMV ncRNA2.7 contains a classical TENT4 binding motif that recruits TENT4 to enable readenylation and mixed tailing (26). This observation provides a potential mechanism by which poly(A) tails on HCMV RNAs may be resistant to CCR4-NOT activity since mixed tails can impede deadenylation (26, 27). To examine this in more detail, we here extend our investigations of poly(A) tail lengths and content on viral RNAs across multiple viral families including human herpesviruses, adenovirus, poxviruses, and coronaviruses. We determine that elongated poly(A) tails are a conserved feature of herpesviruses that is not driven by a mixed-tailing strategy.

## Methods

### Reanalysis of publicly available datasets

Previously published Nanopore datasets generated using the RNA002 DRS methodology for HCMV (25), HSV-1 (28), human adenovirus (29), and the human coronavirus SARS-CoV-2 (30) were downloaded from the Sequence Read Archive (SRA) using the SRA-toolkit (31).

### Monkeypox virus infections

Monkeypox virus (MPXV) strain NL001-2022 was previously isolated at the Erasmus Medical Centre ((32); EvaG Ref-SKU: 010V-04721). MPXV stocks were generated by inoculating 90% confluent Vero cells at a multiplicity of infection (MOI) of 0.01 in Advanced DMEM/F12 (Gibco) supplemented with 10 mM HEPES, 1x GlutaMAX (Gibco) and 1x primocin (Invivogen). The virus was adsorbed for 1 hour 37 °C/5% CO_2_ before washing three times in fresh media. Cells were harvested at day 2 and day 4 (full CPE at day 4) by scraping in advanced DMEM/F12 and frozen. Cell pellets were lysed by freeze-thawing three times and resuspended each time by pipetting. Next, lysates were centrifuged at 2000 x *g* for 5 min and supernatants were stored in aliquots at -80°C before being titrated on Vero cells (32) and used for experiments. For this experiment, two stocks (equal volumes) collected at day 2 and 4 were pooled. Vero cells and normal human dermal fibroblasts (NHDF) were cultured in DMEM (Capricorn Scientific) supplemented with 10% heat-inactivated FBS (Sigma-Aldrich), 2 mM L-glutamine (Gibco) and antibiotics. NHDFs were plated 2 days prior to infection in 6-well plates. Cells were infected with MPXV strain NL001-2022 (passage 4), at an MOI of 1.0 and cultured for 4 or 10 hours. Cells were harvested by removing medium and lysing cells in 1 ml TRIzol per well (Thermo Fisher Scientific). Samples were mixed with 200 μl chloroform (Sigma-Aldrich), centrifuged for 15 min at 12,000 x g at 4°C. RNA was isolated from the aqueous phase using the RNeasy mini kit (Qiagen) followed by DNase treatment using the TURBO DNA-free kit (Ambion). All work with infectious MPXV was performed in a Class II Biosafety Cabinet under BSL-3 conditions.

### Varicella Zoster Virus infections

MeWo cells were maintained in a humidified incubator at 37°C with 5% CO₂ in Dulbecco’s Modified Eagle Medium (DMEM; Gibco #41966-052) supplemented with 8% fetal bovine serum (FBS), 1× glutamine, and 1× penicillin/streptomycin. Cells were passaged at a 1:2.5 to 1:4 ratio two to three times weekly following trypsinization. One day prior to infection, cells were seeded at a 1:2 ratio in 100 mm dishes to achieve approx. 50% confluency at the time of infection. Cell-associated virus (strain EMC-1) was rapidly thawed in a 37°C water bath, diluted in infection medium (DMEM with 2% FCS, 1× glutamine, and 1× penicillin/streptomycin), and added to MeWo cells. After a 2 h incubation at 37°C/5% CO₂, the inoculum was removed, cells were gently washed once with DPBS, and 10 mL of fresh infection medium was added. Four days post-infection, the medium was discarded, cells were rinsed briefly with cold DPBS and scraped into 8 mL of TRIzol. Lysates were transferred to 50 mL conical tubes and stored at -80°C until processing.

### Herpes Simplex Virus Type 2 infections

HSV-2 strain 333 stocks were prepared as previously described (33). ARPE-19 cells were infected at an MOI of 10 and cultured for 10 hours. Cells were harvested by removing medium and lysing cells in 1 ml TRIzol per well. Total RNA was extracted according to the manufacturer’s protocol.

### Kaposi’s sarcoma-associated herpesvirus reactivation

iSLK rKSHV.219 cells (34) were cultured in DMEM supplemented with 10% fetal calf serum, 1% penicillin–streptomycin, 250 mg/mL G418, and 40 μg/mL Puromycin. For reactivation, 2.5×10^6^ iSLK rKSHV.219 cells were seeded in a 10 cm dish. The next day, the lytic cycle was induced by the addition of 1 mM sodium butyrate and 1 µg/ml doxycycline. Cells were harvested at 0 h, 8 h, 24 h, and 72 h post-induction and lysed in TRIzol. Total RNA was extracted according to the manufacturer’s protocol.

### Epstein–Barr virus reactivation

HEK293 cells containing the episomal B95-8-EBV BAC genome (WT^HB9^ from (35) were cultured in RPMI containing 10% fetal calf serum, 1% penicillin–streptomycin, 4 mM L-glutamine and 100 mg/mL hygromycin. For EBV reactivation, 2×10^6^ cells 293-EBV cells were seeded in a 6 cm dish and simultaneously transfected with 3 µg of a 1:1:1 ratio of expression plasmids for BZLF1, BRLF1 and gB (BALF4) mixed with GeneJuice transfection reagent (Millipore) as described previously (36). Forty-eight hours after seeding and transfection, cells were lysed in TRIzol and RNA extracted according to manufacturer’s instructions, but with two additional washes in 70% ethanol before storage at -80°C in 70% ethanol until sequencing.

### Herpesvirus Saimiri (HVS) infection

Owl Monkey kidney (OMK) cells (CRL-1566) were cultured in DMEM cells supplemented with 10% fetal calf serum and 1% penicillin–streptomycin. A BAC-clone of the A11-S4 transformation-deficient strain of herpesvirus saimiri (37) modified to contain a GFP-luciferase expression cassette (38) was propagated in OMK cells and the supernatant harvested and clarified by centrifugation at 500g for 10 minutes. Supernatant was stored in aliquots at -80°C. HVS was titrated by qPCR for the hygromycin gene. 1.25 x 10^6^ OMK cells were seeded in 6cm dish and left overnight before infection with HVS at an MOI of 2500 genomes per cell. RNA harvested 48 hours later as described for EBV.

### Culturing and RNA isolation from human cell lines

All cells were cultured in a humidified incubator, at 37 °C with 5% CO_2_. A549 cells made permissive for SARS-CoV-2 infection by stable expression of human ACE2 (30) were maintained in DMEM, 10% fetal bovine serum, and penicillin/streptomycin. MeWos were cultured in DMEM supplemented with 8% FBS, 1% Pen/Strep, 1% L-glutamine, while NHDFs (passage 12) were cultured in DMEM 10%FCS, 1%P/S, and 1% L-glutamine. ARPE-19 cells were cultured in DMEM/Nutrient mixture F-12 Ham medium (Sigma-Aldrich), 8% FCS, 1% penicillin/streptomycin, 2 mM L-Glu. In all cases, cells at 90% confluence were harvested by adding TRIzol and RNA extracted according to manufacturer’s protocol with addition of GlycoBlue (ThermoFisher) during the first precipitation step.

### Direct RNA Sequencing

Poly(A) RNA was isolated from total RNA using Dynabeads (Invitrogen) with 133 µl beads added to 25 µg of total RNA. Nanopore DRS libraries were prepared according to the standard SQK-RNA002 protocol or the Deeplexicon multiplexing protocol (39) and sequenced for 24-48 hours on R 9.4.1 flowcells using a MinION Mk.1b.

### Reference genomes and transcriptomes

The human genome assembly (GRCh38.p14) was obtained from Ensembl (https://www.ensembl.org/index.html) while the human transcriptome (v47) was obtained from Gencode (https://www.gencodegenes.org/human/). Viral reference genomes downloaded from Genbank (https://www.ncbi.nlm.nih.gov/genbank/) were obtained with the following accession numbers: VZV Dumas (NC_001348.1), HSV-1 KOS (KT899744.1), HSV-2 333 (LS480640.1), HCMV TB40/E (EF999921.1), KSHV GK18 (NC_009333.1), HAdV-F41 (ON561778.1), Mpox UK1 (MT903343.1), and SARS-CoV-2 (MN985325.1). The HVS-A11-S4 BAC sequence is modified from the HVS-A11 sequence (NC_001350) to include the Sac I deletion of transforming genes and insertion of the BAC-associated sequence. The EBV B95-8-BAC sequence is modified from accession number V01555 to reduce the number of IR1 repeats from 11 to 6, and to add the BAC sequence. The transcriptome annotations for HSV-1 KOS and HCMV TB40/E were derived from custom annotations that are available from https://github.com/DepledgeLab/polyAtails. The EBV transcriptome annotation (40) was obtained from ebv.org.uk.

### Data parsing, alignment, and estimation of poly(A) tail lengths

For all DRS datasets, high-accuracy basecalling was performed with Guppy v6.1.7 [ *-c rna_r9.4.1_70bps_hac.cfg -r --calib_detect --trim_strategy rna --reverse_sequence true -x auto*]. Where the basecalling outputs were analysed with *ninetails,* we additionally included the writing to fast5 step [ *--fast5_out*]. Resulting fastq files were aligned against the respective reference genome [ *-ax splice -k14 -uf --secondary=no*] and/or transcriptome [ *-ax map-ont-L -p 0.99 -uf --secondary=no*] using minimap2 (41) and parsed to generate sorted BAM files using SAMtools v1.15 (42) in which only primary alignments were retained. Estimation of poly(A) tail lengths was performed using nanopolish v0.14 (21).

### Non-adenosine nucleotide detection in poly(A) tails using ninetails

Selected DRS datasets have been examined for non-A nucleotides in poly(A) tails with ninetails 1.0.3(24) utilising the *check_tails* wrapper function [ *pass_only=FALSE*] using nanopolish poly(A) tail length estimation and the Guppy sequencing summary. Predictions were reclassified to increase precision with *reclassify_ninetails_data* [ *ref=’hsapiens’*], merged with *merge_nonA_tables* [ *pass_only=FALSE*] and separately collapsed by transcript with *summarize_nonA*.

### Generation of R plots & R packages used

All plotting was performed using Rstudio (https://posit.co/download/rstudio-desktop/) with R v4.4.3 and the following packages: data.table (https://r-datatable.com), ggplot2 v3.5.2 (43), dplyr v1.1.4 (https://dplyr.tidyverse.org/), tidyverse (44), tidyr (https://tidyr.tidyverse.org/), stringr v1.5.1 (https://stringr.tidyverse.org/), ggpubr v0.6.0 (https://rpkgs.datanovia.com/ggpubr/), readxl (https://readxl.tidyverse.org/), glmmTMB v1.1.11 (https://github.com/glmmTMB/glmmTMB), forcats v1.0.0 (https://forcats.tidyverse.org/), reticulate v1.42 (https://rstudio.github.io/reticulate/), viridis v0.6.5 (https://sjmgarnier.github.io/viridis/), cowplot v1.1.3 (https://wilkelab.org/cowplot/),rstatix v0.7.2 (https://rpkgs.datanovia.com/rstatix/), zoo v1.8 (https://zoo.r-forge.r-project.org/),ggthemes v5.1.0 (https://github.com/jrnold/ggthemes), FSA v0.9.6 (https://fishr-core-team.github.io/FSA/), ggsignif v0.6.4 (https://osf.io/preprints/psyarxiv/7awm6), patchwork v1.3.0 (https://patchwork.data-imaginist.com), and ninetails v1.0.3 (24).

### Data availability

All R scripts and input datasets are available in the https://github.com/DepledgeLab/polyAtails repository. Raw DRS002 fast5 (nanopore) datasets generated as part of this study are available at the ENA/SRA under the accession number PRJEB98222 with the exception of the HVS and EBV datasets which are available via accession numbers PRJEB104941 and PRJEB105217, respectively.

## Results

### Extended poly(A) tails are a shared feature of herpesvirus mRNAs

We performed nanopore DRS on poly(A)+ RNA isolated from three human cell lines commonly used to support virus infection: A549 lung adenocarcinoma cells, MeWo skin melanoma cells and normal human dermal fibroblasts (NHDFs). Following alignment to the human transcriptome and filtering to retain only protein-coding mRNAs, we generated violin plots to summarise the global distribution of poly(A) tail lengths (Fig. 1a). These data demonstrate that while poly(A) tail lengths on individual RNAs vary significantly, the majority cluster around modal values of 48 – 54 nt (Table S1) across the three cell types tested, in agreement with previous studies using both nanopore and orthogonal methods of tail measurement in other human cell lines (10, 11, 19, 45). The width of violin plots relates the density of data and thus allows for interpretation not only of the modal value but also the proportion of poly(A) tail lengths that fit the modal value i.e., a slightly larger proportion of mRNAs in A549 cells have poly(A) tail lengths of ∼52 nt when compared to NHDFs. Using a combination of new and existing nanopore DRS datasets (all SQK-RNA002 chemistry), we examined the poly(A) tail length distributions, separately, for host and viral protein-coding mRNAs from productively infected cells with six different human herpesviruses (Fig. 1b-g) including HSV-1 strain KOS-infected NHDFs, HSV-2 strain 333-infected ARPE-19 cells, VZV strain EMC-1 infected MeWos, HCMV strain TB40/E-infected NHDFs, and reactivated KSHV iSLK.219 and EBV-B-95-8 BAC HEK293 cells. In addition, we examined productive infection by a non-human primate gammaherpesvirus, Herpesvirus Saimiri (HVS), in owl monkey kidney (OMK) cells (Fig. 1h). Each was profiled at a late-stage of productive infection, using high MOIs where possible (i.e. HSV-1, HSV-2, and HCMV) to maximise the relative proportion of viral reads (Fig. 1i). Our analysis demonstrated that while poly(A) tail length distributions on human mRNAs did not significantly differ between uninfected cells and cells infected with HSV-1, HSV-2, HVS, and the reactivated KSHV iSLK and EBV HEK293 cells (modal values of 42 – 52, Table S1), we observed notably increased poly(A) tail lengths on human mRNAs for VZV and HCMV infected cells (modal values of 80 and 94, respectively, Table S1). For herpesviral protein-coding mRNAs, we observed uniformly longer poly(A) tail length distributions (modal values of 96 – 150, Table S1). Among the herpesvirus tail distributions, VZV tails were the longest, with a modal length of 150 nt and HSV-2 tails the shortest (96 nt modal length), while still markedly exceeding those of host mRNAs. Notably, the shapes of the distributions also differed e.g. for HSV-2 the density of poly(A) tail lengths is more widely distributed, resulting in a narrower plot. Together, these data demonstrate that longer poly(A) tails represent a shared feature of alpha, beta and gammaherpesvirus mRNAs, in both reactivation and productive replication contexts, which could contribute to efficient viral gene expression and represent a novel viral tactic to subvert the host.

**Figure 1:**
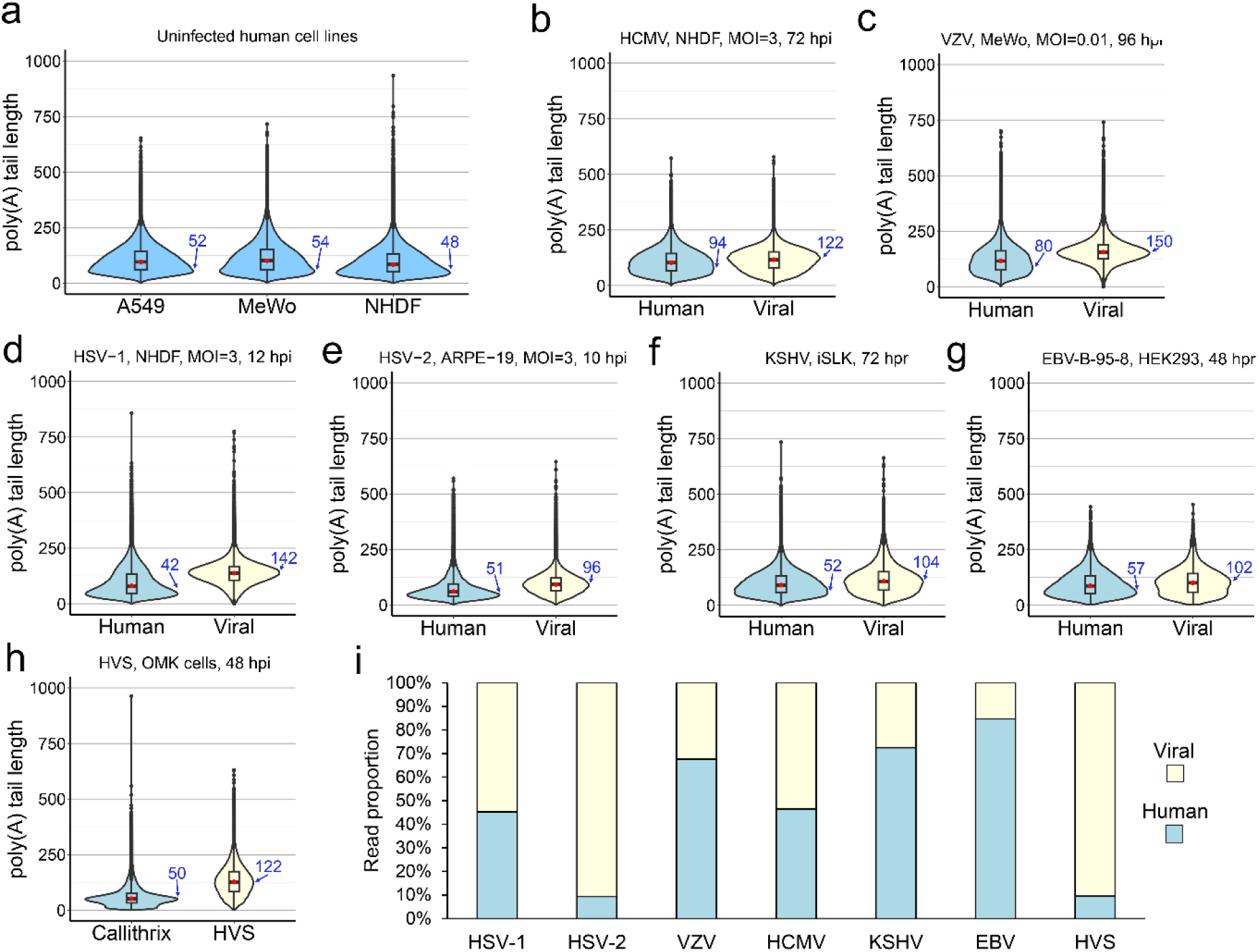
Extended poly(A) tails are a shared feature of herpesvirus mRNAs. Violin plots denoting poly(A) tail length distributions for protein coding mRNA populations of **(a)** three uninfected human cell lines, **(b)** HCMV, **(c)** VZV, **(d)** HSV-1, **(e)** HSV-2, **(f)** KSHV, **(g)** EBV, and **(h)** HVS. **(i)** Relative proportions of human (blue) and viral (yellow) reads in each dataset are shown. Modal poly(A) tail lengths for each dataset are written in blue. hpi – hours post infection, hpr – hours post reactivation.

### Elongated poly(A) tails do not represent a uniform feature of viral infection

We next considered whether longer poly(A) tails could reflect a broader viral strategy to maximize viral gene expression. We thus extended our analysis to examine poly(A) tail length distributions (Fig. 2a-d) on human and viral protein-coding mRNAs for a positive sense RNA virus that replicates in the cytoplasm, SARS-CoV-2, a cytoplasmic-replicating DNA virus, Mpox virus (MPXV; Clade 1b, strain NL1), and a nuclear-replicating DNA virus from a different virus family, human Adenovirus type 41 (hAdV41). We further examined multiple timepoints for the hAdV41 (12, 24, 48 hours post infection, hpi) and Mpox (4, 10 hpi) infected cells (Fig. 2b-c). Similar to VZV and HCMV infections but to a greater extent, SARS-CoV-2 infection led to an increase in human mRNA poly(A) tail lengths (modal value of 170) and a broadening of the overall tail distribution. By contrast, hAdV41 and Mpox infections did not impact on poly(A) tail length distributions on human protein-coding mRNAs (modal values of 46-50, Table S1).

**Figure 2:**
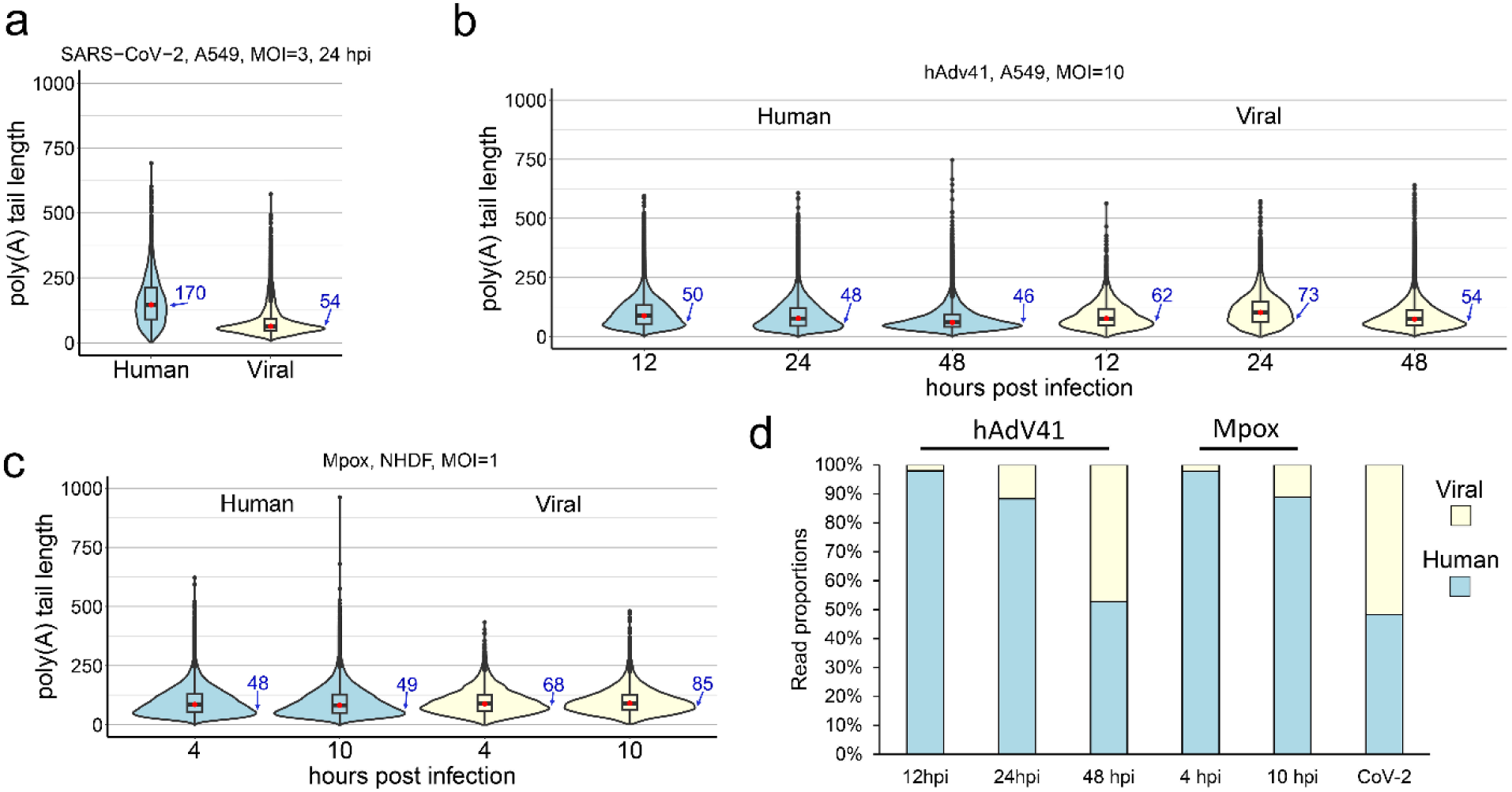
Long poly(A) tails is not a universal viral strategy. Violin plots denoting poly(A) tail length distributions for protein coding mRNA populations of **(a)** SARS-CoV-2 at 24 hpi, **(b)** Human Adenovirus 41 at 12, 24, and 48 hpi, and **(c)** Mpox virus at 4 and 10 hpi. (d) Relative proportions of human (blue) and viral (yellow) reads in each dataset are shown. Modal poly(A) tail lengths for each dataset are written in blue.

Replication of coronaviruses produces polyadenylated genomic RNAs (gRNAs) and, through discontinuous transcription, larger amounts of subgenomic mRNAs (sgmRNAs) that encode structural and accessory proteins. In our analyses of SARS-CoV-2 RNAs, nanopolish could not accurately identify the poly(A) tails on the few gRNA reads present, whereas poly(A) tail lengths could be identified for the majority of sgRNA reads. The poly(A) tail lengths for viral sgmRNAs were tightly distributed around ∼50 nt, consistent with those previously reported (46). Adenoviral mRNAs exhibited a lifecycle-dependent poly(A) tail length advantage over host mRNAs (Fig. 2b, Table S1). At 12 hpi, the distribution of viral poly(A) tails skewed slightly longer than host tails (modal length 62 vs 50 nt). This increased further at 24 hpi (modal length 73 vs 48) but decreased again by 48 hpi (54 vs 46 nt). Consistently, a similar pattern of shorter viral poly(A) tail lengths as infection progressed was previously reported for adenovirus type 5 (47). It is notable that although, like herpesvirus RNAs, adenoviral RNAs are transcribed and polyadenylated by the host transcriptional machinery they do not display similarly long modal poly(A) tail lengths. Poxviral mRNAs are generated in the cytoplasm and poly(A) tails are added by a viral poly(A) polymerase (2). We observed Mpox poly(A) tail distributions were also slightly longer than those on host mRNAs with a modal length of 68 - 85 nt (Fig. 2c, Table S1). While not as long as those of herpesvirus mRNAs, these extended tails could benefit the translation of poxvirus mRNAs. Indeed, PABPC1 is enriched at poxvirus replication factories where viral translation occurs, and enhanced eIF4F translation initiation complex formation is detected in infected cells (48, 49).

The human transcriptome comprises a mixed population of mRNAs at various ages post-synthesis and stages of decay. By contrast, the coordinated surge of viral gene transcription upon infection with herpes- and adenoviruses is expected to yield a relatively “young” viral transcript pool. However, since poly(A) tails on herpesviral mRNAs are much longer than for other viruses – even the biologically similar adenovirus – it is unlikely that mRNA age alone can fully explain the extended poly(A) tail lengths of herpesviral mRNAs. Contrasting tail lengths on viral mRNAs in coronavirus and poxvirus infections underline that extended tails are not a universal feature of viral RNAs and that effects on host mRNA poly(A) tails vary.

### Long poly(A) tails also occur on herpesviral noncoding RNAs

The major characterised eukaryotic deadenylation complexes PAN2-PAN3 and CCR4-NOT are predominantly cytoplasmic, with the activity of the latter being linked to translation (1). To investigate the mechanism behind long poly(A) tails on herpesviral mRNAs, we also examined herpesviral ncRNAs. A prior report demonstrated that HCMV ncRNAs encoded in the Toledo strain contain poly(A) tails of differing lengths (50). Here, we observed a similar result, extending our observation of poly(A) tail lengths on HCMV TB40/E strain protein coding mRNAs to also include the HCMV encoded nuclear-retained RNA4.9, cytoplasmic RNA2.7 and nuclear-cytoplasmic RNA1.2, and contrasted these with a human nuclear-retained paraspeckle ncRNA (NEAT1) (Fig. 3a). We found both RNA4.9 and RNA2.7 to contain longer poly(A) tails (modal values of 152 and 156) than HCMV mRNAs (modal value 122). By contrast, RNA1.2 (modal value 95) showed a shorter poly(A) tail length distribution when compared to HCMV mRNAs. Of note, the poly(A) tails on HCMV RNAs of all types remain significantly shorter than on NEAT1 (modal value 269). We next performed a similar analysis for KSHV, contrasting the poly(A) tail length distributions for the well-characterised nuclear-retained PAN ncRNA (51) with that of the Kaposin mRNA, recently characterised to partially localise to and scaffold nuclear speckles (52), and all other KSHV mRNAs (Fig. 3b). Surprisingly, we observed the poly(A) tail length distribution and modal value of PAN (69 nt) to be similar to that of cellular mRNAs and quite different from the KSHV mRNA population (104 nt). Kaposin mRNA did not display an obviously different tail length distribution from the remaining KSHV mRNAs. Together these data demonstrate that poly(A) tail lengths vary dramatically across different viral ncRNAs, implicating transcript-specific mechanisms of control that can augment the generalised extended poly(A) tails on herpesviral RNAs. Moreover, herpes RNA tail lengths are not singularly defined by nuclear retention or translation status.

**Figure 3:**
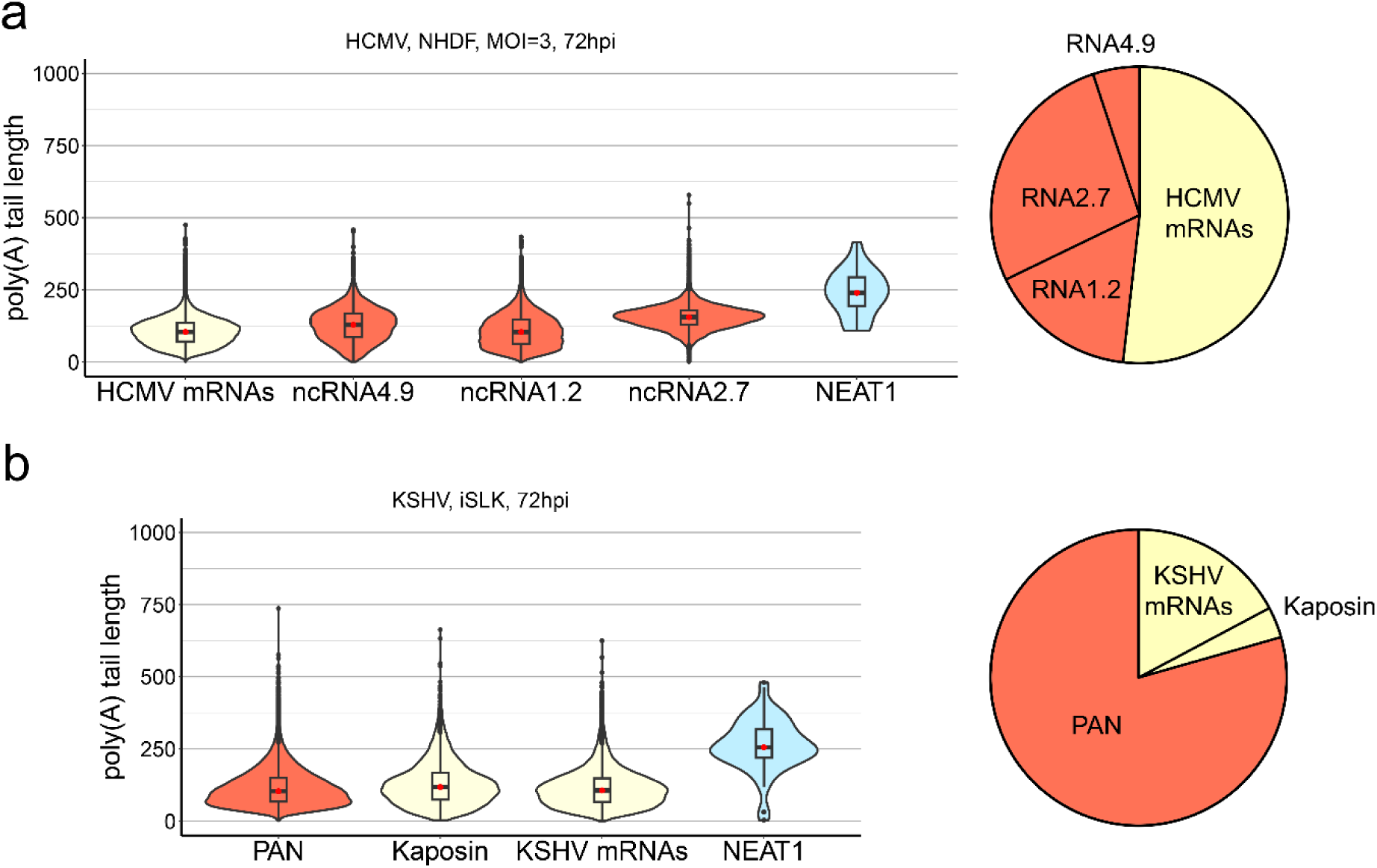
Long poly(A) tails on herpesviral ncRNAs are not associated with nuclear retention. **(a)** Violin plots denoting poly(A) tail length distributions for all HCMV mRNAs, three polyadenylated HCMV ncRNAs; RNA4.9 (nuclear restricted), RNA1.2 (nucleocytoplasmic), and RNA2.9 (cytoplasmic), contrasted against the cellular ncRNA NEAT1_1. The pie chart denotes the relative proportions of HCMV mRNAs, RNA1.2, RNA2.7, and RNA4.9**. (b)** Violin plots denoting poly(A) tail length distributions for the KSHV ncRNA PAN, the KSHV kaposin mRNA, and all remaining KSHV mRNAs, contrasted against the cellular ncRNA NEAT1_1. The pie chart denotes the relative proportions of PAN, Kaposin, and all other KSHV mRNAs.

### HSV-1 and HCMV encoded RNAs display differential tail length patterns during infection

To better understand the kinetics of adenylation/deadenylation during infection, we next performed an analysis of poly(A) tail length differences on HSV-1 strain KOS mRNAs sampled at 3, 6, and 12 hpi (Fig. 4, Table S2). A global analysis of viral poly(A) tail length distributions at each time point demonstrated that the overall distribution remained broadly similar with only a slight decrease in modal values during infection (3h – 148 nt, 6h – 142 nt, 12h – 142 nt, Fig. 4a). To improve the resolution of our analysis, we subsequently realigned our nanopore datasets against a high-resolution annotation of the HSV-1 strain KOS transcriptome. This enabled us to analyse poly(A) tail lengths for each mRNA individually, measuring changes in the median poly(A) tail length over time (Fig. 4b). Importantly, this demonstrated that while most median poly(A) tail lengths on viral mRNAs remained similar or decreased over infection, three transcripts, US12 (ICP47), RS1 (ICP4) and RL2 (ICP0), showed evidence of median poly(A) tail lengths increasing by ≥ 27 nt between at least two timepoints (Fig. 4c, d). A 27 nt increase could accommodate an additional PABPC molecule (14) and thus affect mRNA stability and, or, translation. To determine whether transcription kinetics were linked to viral poly(A) tail status, we grouped transcripts according to their temporal designation (*Immediate-Early*; *Early*; *Late*; *Leaky Late*; *True Late*) (Fig. 4e). Here we observed that *Immediate-Early* transcripts were outliers, with those detected at 6 and 12 hpi having longer tails than at 3 hpi, with the exception of UL54 (ICP27) for which poly(A) tail lengths do not change during infection. By contrast, *Early* transcripts generally had shorter tails at 6 and 12 hpi compared to 3 hpi. This pattern was also observed for *Late* transcripts and, most obviously, for *True Late* transcripts, which displayed a 9 nt reduction in median poly(A) tail lengths from 6 to 12 hpi. This suggests that though their tail lengths are long, most HSV-1 transcripts are not completely refractory to deadenylation. Exceptions to this trend highlight that transcript-specific regulation of poly(A) tail length may additionally be involved. Among those that show increased tail lengths at later stages of infection, RL2 (ICP0) mRNA has previously been reported to display nuclear retention (53). This partial segregation from the cellular cytoplasmic deadenylation machinery may thus explain this sustained increase in poly(A) length.

**Figure 4:**
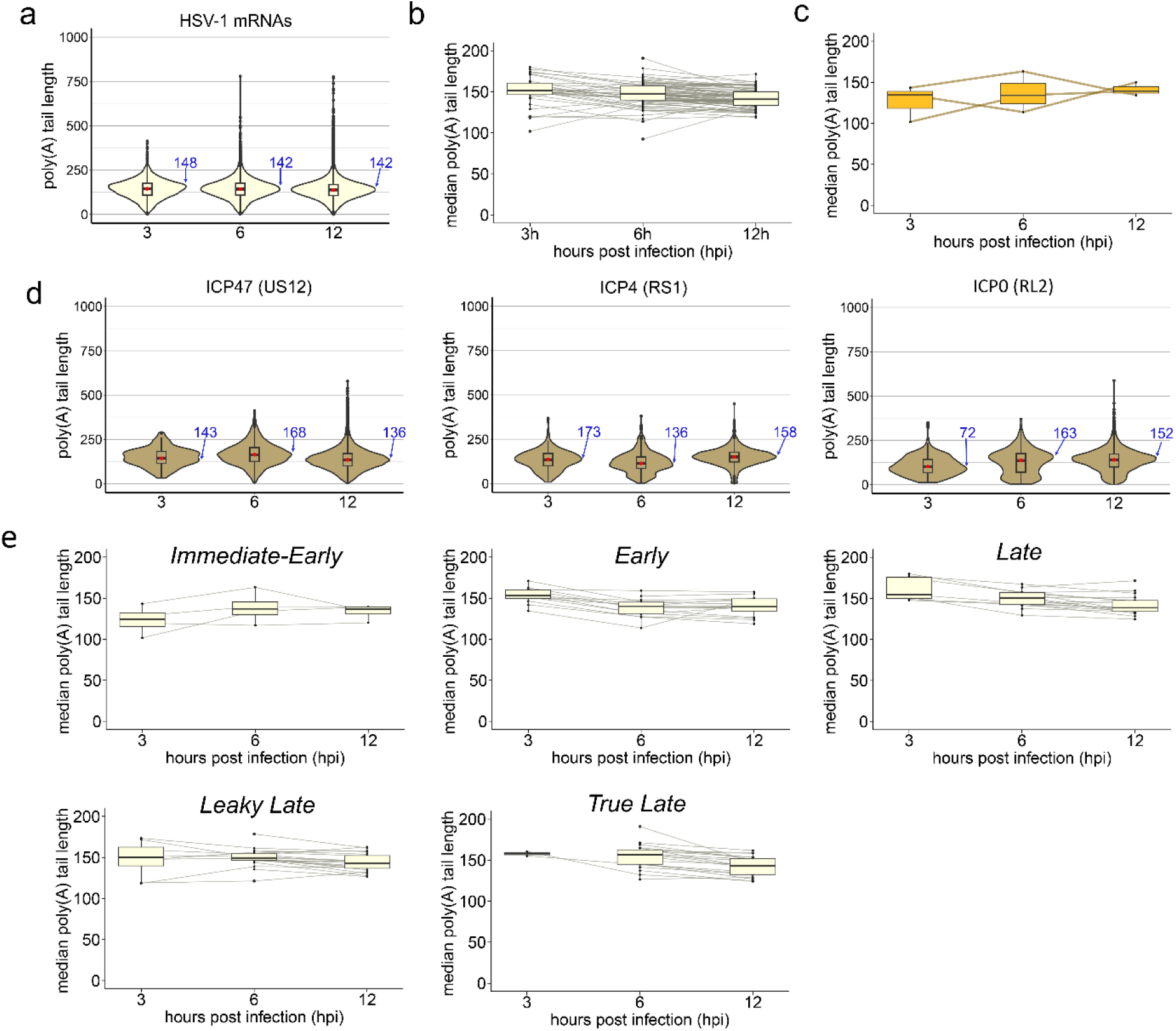
HSV-1 RNAs display differential adenylation patterns during infection. **(a)** Violin plots denoting HSV-1 mRNA poly(A) tail length distributions across timepoints. **(b)** Boxplots denoting changes in median poly(A) tail length for each HSV-1 mRNA during infection. **(c)** As **(b)** but showing only the subset of HSV-1 mRNAs that display increased tail lengths over the course of infection. **(d)** Violin plots denoting poly(A) tail length distributions for representative examples of HSV-1 mRNAs that show increased of > 25 nt in median poly(A) tail lengths between at least two time-points in infection. **(e)** Boxplots denoting changes in median poly(A) tail length for HSV-1 mRNA, separated according to kinetic class: *alpha* – *Immediate-Early*, *beta* – *Early*, *gamma* – *Late*, *gamma 1* – *Leaky Late*, *gamma 2* – *True Late*. Modal poly(A) tail lengths for each dataset in (a) and (d) are written in blue.

To determine whether there is also evidence of transcript-specific poly(A) tail length changes across HCMV RNAs we analysed HCMV strain TB40/E mRNAs expressed at 24-, 48-, and 72-hours post-infection (Fig. 5, Table S3). Similar to HSV-1, we observed that poly(A) tail lengths on HCMV mRNAs decrease only slightly over the course of infection, while a small subset shows more dramatic increases and/or decreases (Fig. 5a). By contrast, all four polyadenylated HCMV ncRNAs displayed very long tails (medians of 149 – 219 nt) at 24 hpi which decreased sharply to 108 – 162 nt by 48 hpi and remained relatively unchanged (104 – 156 nt) at 72 hpi (Fig. 5b). An analysis of HCMV mRNAs by temporal class (as defined in (54)) revealed a similar pattern to the whole population of small decreases in long poly(A) tail lengths over the viral lifecycle. Within these classes, as with HSV-1, individual outliers were identified, such as UL17-1 and US24-1 in temporal class 3 and US32-1 in temporal class 7, whose tails increased between 24 and 48hpi (Fig. 5c). These data offer further support that herpesviral RNAs can be deadenylated, and we find no evidence that the timing of their synthesis explains their extended length.

**Figure 5:**
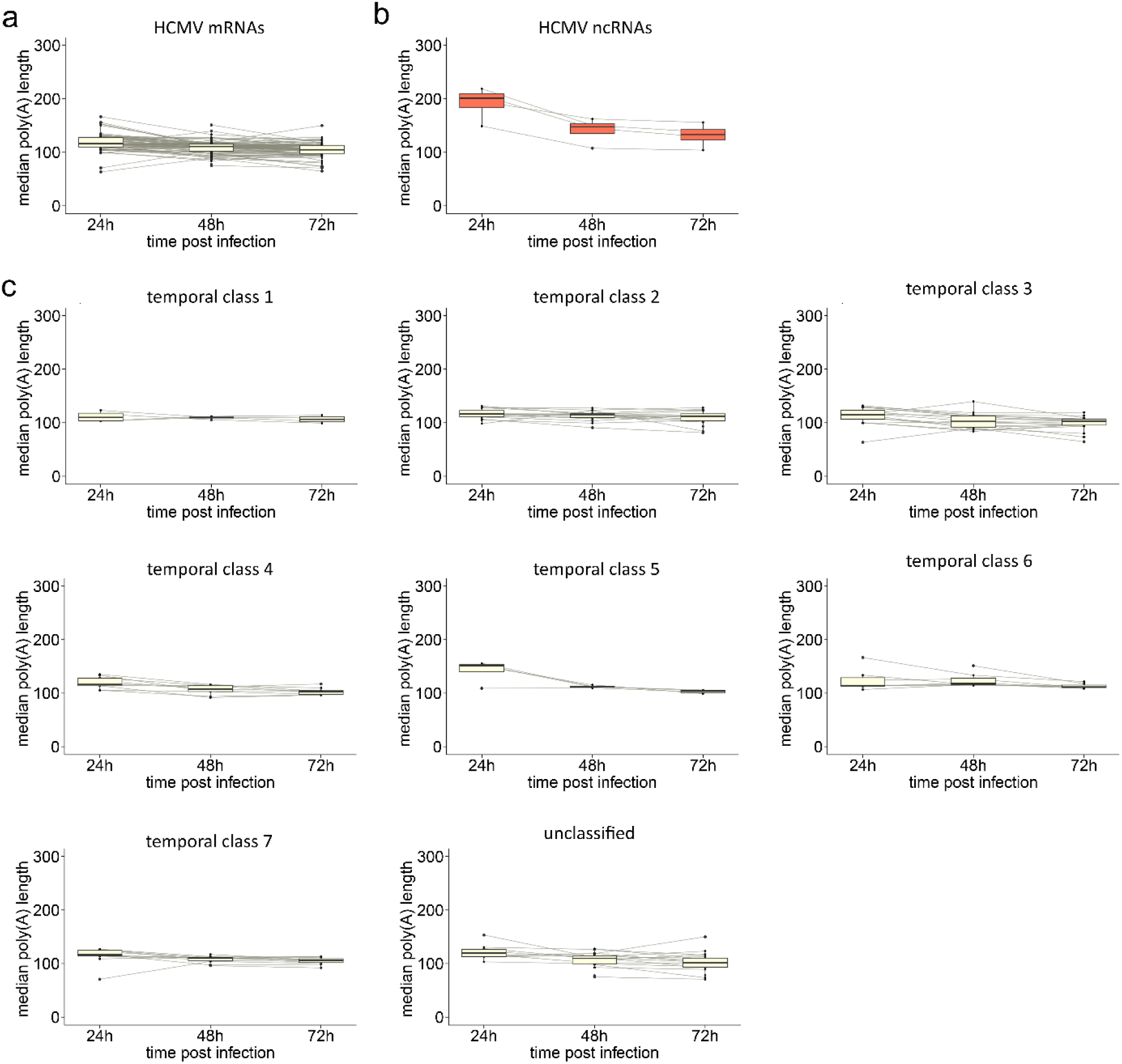
HCMV mRNAs generally display consistent poly(A) tail lengths during infection. **(a-b)** Boxplots denoting changes in median poly(A) tail length for each HCMV **(a)** mRNA and **(b)** ncRNA during infection. **(c)** Boxplots denoting changes in median poly(A) tail length for HCMV mRNAs, separated according to seven kinetic classes defined by (54).

### Increased rates of mixed-tailing during herpesvirus infections does not explain longer tails on viral mRNAs

Incorporation or addition of non-adenosine nucleotides to poly(A) tails by non-canonical PAPs, known as ‘mixed tailing’, can impede mRNA deadenylation (55). A recent study identified a motif in HCMV RNA2.7 capable of recruiting such a PAP and using TAIL-seq found non-As within HCMV poly(A) tails (26). To examine the possibility that herpesviral poly(A) tails are generally extended due to mixed tailing, we applied *ninetails* (24) to DRS datasets to quantify non-adenosine nucleotide incorporation events. Using an *in vitro* transcribed (IVT) RNA with a pure poly(A) tail, we established that *ninetails* performs well with a low false-positive rate of around 1% and a bias toward false-positive incorporations of G (Fig. 6a, Fig. S1a). In uninfected NHDFs we detected mixed tails on 6.5% of transcripts (Fig. 6a), which is below that found on HeLa cell transcripts by orthogonal methods (17% by PAIso-seq (56); 12.5% by FLAM-seq (45)) but closer to that of a prior *ninetails* analysis (11%, (24)). Exclusion of the extreme (∼5 – 10 nt) 3’ terminal nucleotides from *ninetails* analysis could account for this difference (24). Consistent with previous FLAM-seq and *ninetails* studies, we also observe that C incorporations are more common than U and G (Fig. 6b, Fig. S1a). Remarkably, we found that HCMV-infection of NHDFs leads to an increase in mixed tail frequencies on cellular mRNAs over the course of infection, up to 13.3% by 72hpi (Fig. 6a) with a proportional decrease in G and C incorporations and increase in U incorporations (Fig. 6b). For HCMV mRNAs, mixed-tailing rates increased from 6.6% to 12.2% with infection time and consistently exceeded that of human RNAs (Fig. 6c, Fig. S1b), albeit while maintaining similar distributions of C, U, and G incorporations (Fig. 6d). This suggests that HCMV infection upregulates a mixed tailing mechanism/s during infection that is not selective for viral over host mRNAs. When we examined additional herpesvirus datasets, intriguingly we observed particularly high rates of mixed tailing on viral transcripts in late-stage HSV-2 infected ARPE-19s (23%) but lower rates for both KSHV (6.1%) and VZV (7.1%) (Fig. 6c), suggestive of differential ability of herpes viruses to coordinate this effect. Mixed tail frequency and poly(A) tail length distribution are therefore not correlated during HCMV infection or among different herpesviruses. Since the frequency of mixed tailing on herpesvirus mRNAs is at its maximum less than 25%, these data demonstrate that mixed tailing cannot account for the consistently extended poly(A) tails of herpesvirus RNAs.

**Figure 6:**
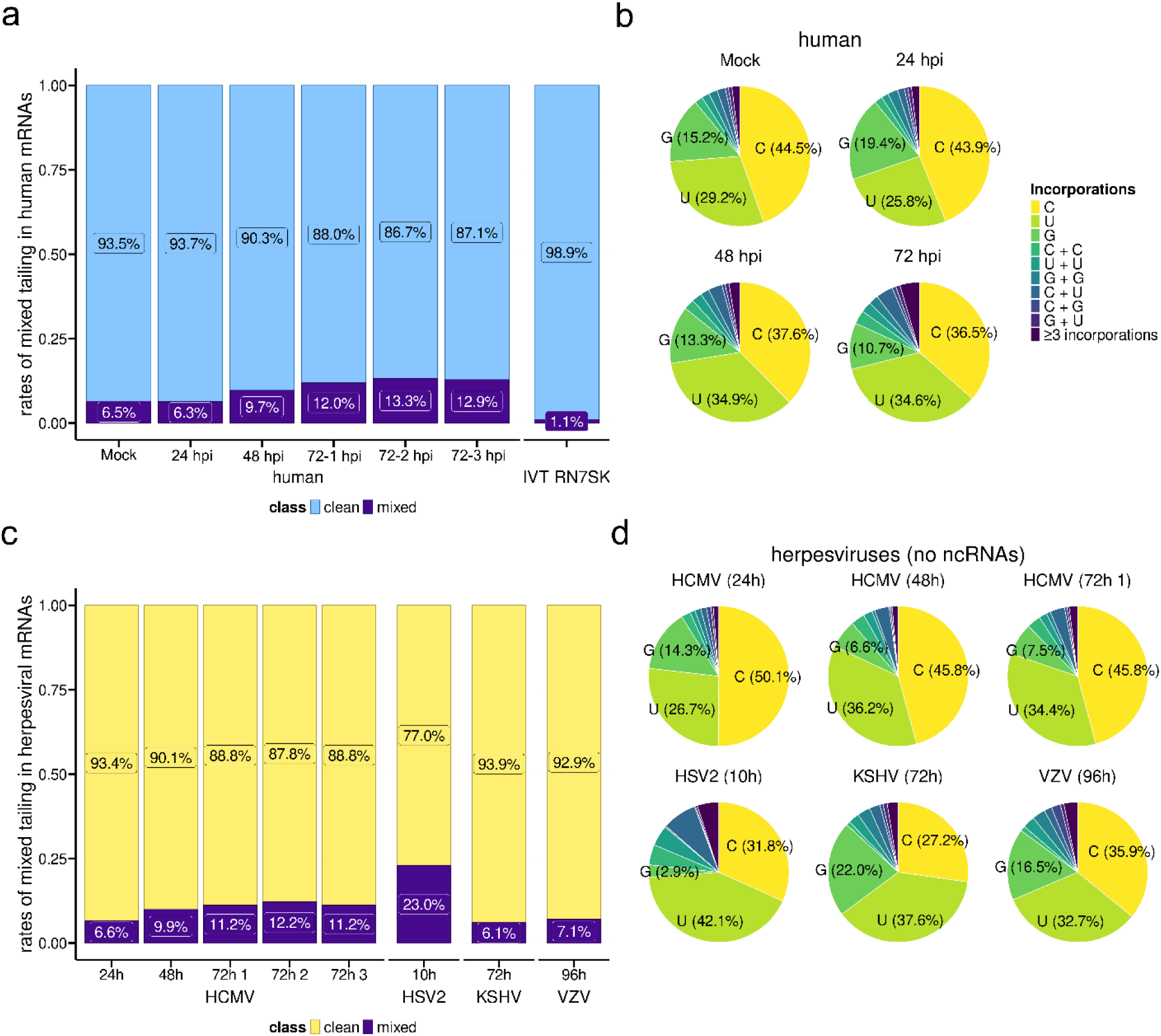
Incorporation of non-adenosine nucleotides in human and herpesviral mRNA poly(A) tails. **(a)** Incidences of mixed-tailing (i.e. non-adenosine nucleotide incorporations) predicted by *ninetails* (24) for cellular mRNAs obtained from mock and HCMV-infected cells at the indicated timepoints. A synthetic RN7SK RNA with a clean tail is also included. **(b)** Breakdown of nucleotides present in mixed tails of cellular mRNAs by timepoint. **(c)** Incidences of mixed-tailing in herpesviral mRNAs obtained from infected cells at the indicated timepoints **(d)** Breakdown of nucleotides present in mixed tails of herpesviral mRNAs by timepoint.

### Mixed-tailing on individual viral and host transcripts

We next examined the rates of U, G, and C incorporations across HCMV RNAs and human mRNAs supported by at least 30 reads, as well as highlighting the most abundant transcripts in the human (n=50) and HCMV datasets (n=82) (Fig. 7a-c). We determined that HCMV infection induces a generalized increase in mixed tailing rates across both viral and cellular mRNAs (Fig. 7a-c, Fig. S2). Focusing specifically on guanine incorporation, we validated the prior observation (26) that HCMV RNA2.7 displays comparatively high G-incorporation rates in poly(A) tails (Fig. 7b), consistent with the activity of the interacting TENT4. Surprisingly, we also observed similarly high G-incorporation rates for UL69 which encodes a multi-functional protein that is homologous to HSV ICP27 and conserved across all human herpesviruses (57) suggesting it may also be a substrate for TENT4. Similarly, several human mRNAs also displayed increased G-incorporation rates following HCMV infection including B2M-211 and HSP9011A-201 (Table S4). Notably, examples of highly expressed HCMV mRNAs with G-incorporation rates that did not change during the infection period (UL132) or that showed significantly decreased incorporation rates (UL22A) were also observed, demonstrating that viral transcript specific features can also influence G-incorporation in poly(A) tails. Full details of all human and viral mRNAs examined and their relative C- G- and U-incorporation rates over time are shown in Table S4. Finally, we examined the relative location of C-, G-, and U-incorporations within poly(A) tails on human and HCMV mRNAs and observed that for HCMV RNAs (RNA2.7, UL69) and human mRNAs (B2M-211 and HSP9011A-201) with high G-incorporation rates, there was a significant enrichment of G-incorporations toward the 3’ end of poly(A) tails, consistent with the addition of G nucleotides during cytoplasmic extension of poly(A) tails and their resistance to 3’ decay (Fig. 7d. Fig. S1c). In contrast C- and U-incorporations were more evenly distributed through the tails on human mRNAs, and HCMV RNA2.7 early in infection. At 72 hpi however C- and U-incorporation in the RNA2.7 poly(A) tail were also enriched toward the 3’ end. Thus, our data supports the contribution of mixed-tailing to the extended poly(A) tail length of non-coding HCMV RNA2.7 and identifies additional viral and host mRNAs which may be regulated in this manner.

**Figure 7:**
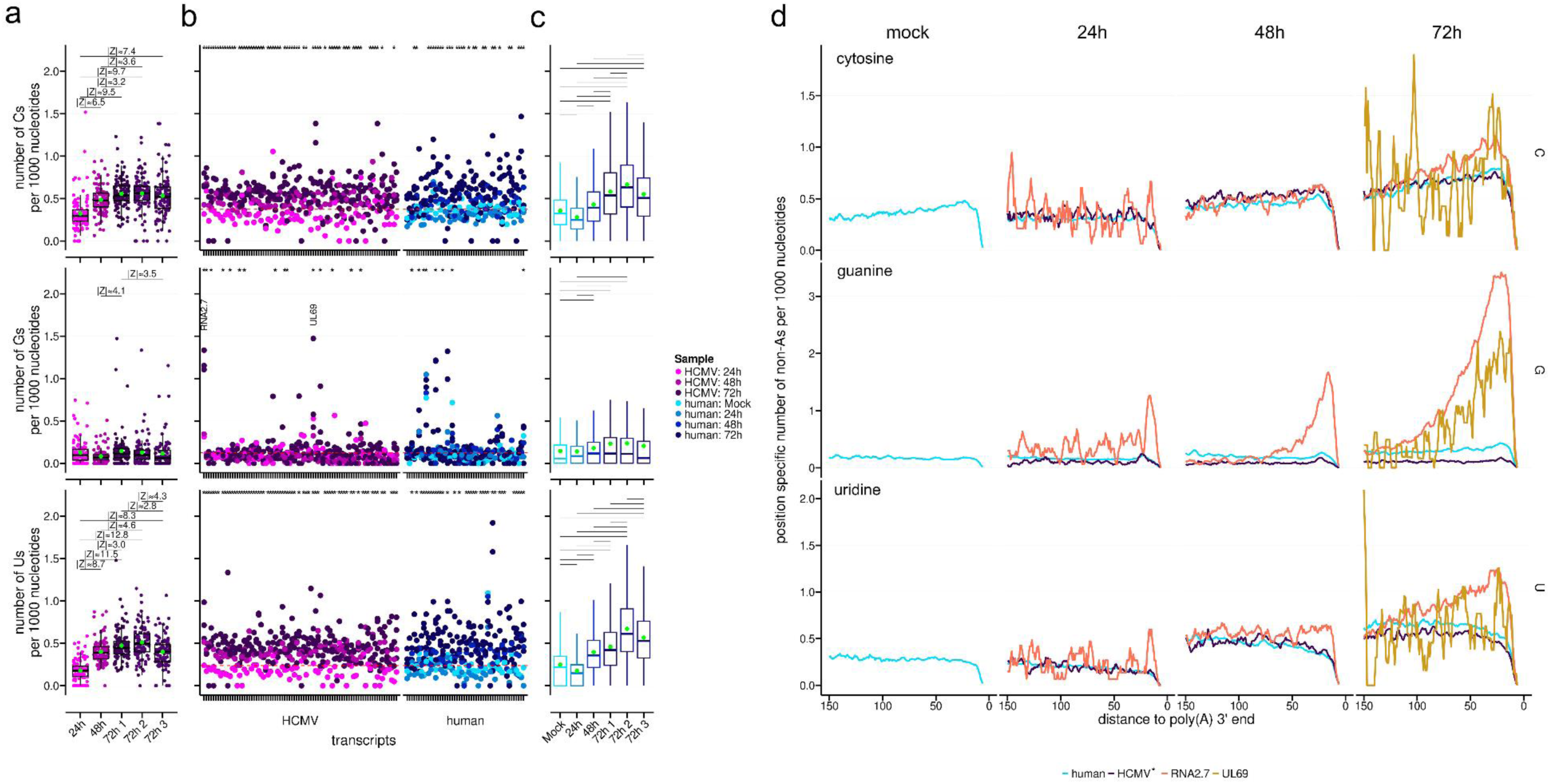
Transcript- and nucleotide-level incorporation of non-adenosine nucleotides in human and HCMV RNA poly(A) tails. **(a)** Frequency of specific non-As at the per nucleotide level by HCMV RNAs (coverage ≥ 30 reads) across different timepoints, as predicted by *ninetails* (24). Additional 72h biological replicates are shown in Figure S1. Boxplots whiskers depict 1.5 * IQR with the mean is shown by the green dot. A Kruskal-Wallis rank sum test was applied on all groups for each non-A nucleotides, followed by Dunn’s test with Bonferroni correction. Absolute Z-scores with significant adjusted p-values (≤0.05) are shown above the boxplots. **(b)** Frequency of C, G, and U incorporations in poly(A) tails at the per nucleotide level for selected HCMV and human mRNAs across three infection timepoints. For the HCMV panel, pooling the top 50 mRNAs from each time point produced 82 unique transcripts. The human panel shows the 50 highest-coverage mRNAs in the mock sample (red dashed lines = mean frequencies). The red dashed line indicates the mean frequency of the relevant incorporation in the mock-infected dataset. A negative-binomial generalized linear mixed model was fitted to the non-A frequencies comparing 72h vs mock-infection. P-values were adjusted via the Benjamini-Hochberg procedure. Transcripts with significant adjusted p-values (≤0.05) are marked with an asterisk.**(c)** Frequency of C, G, and U incorporation per 1000 nucleotides within poly(A) tails of human mRNAs (coverage ≥ 30 reads) across different timepoints. Boxplots whiskers depict 1.5 * IQR, the mean is shown by the green dot. Kruskal-Wallis rank sum tests were applied on all groups for each of the three non-adenosine residues followed by Dunn’s test with Bonferroni correction. Significant adjusted p-values (≤0.05) are marked above the boxplots, complete Z-scores and adjusted p-values can be found in Table S5. **(d)** Position specific frequency of non-A nucleotides in the poly(A) tail of different transcripts and timepoints, defined by their distance of the terminal 3’ A nucleotide. Human includes all reads of human mRNAs. HCMV includes all reads from HCMV RNAs except RNA2.7 and UL69, which are shown separately. Plotted frequencies are calculated by the number of specific non-adenosine residues with a certain distance *d* to the 3’ end relative to the total number of reads with a poly(A)tail length ≥ *d*, averaged with a sliding window of 5 nucleotides.

## Discussion

In this study, we have shown that poly(A) tail lengths on herpesviral RNAs are significantly longer than those on both human protein-coding mRNAs and a diverse set of other DNA and RNA viruses. While the functional consequence of this pan-herpesviral strategy remains untested, we can hypothesise that mRNAs with longer tails are associated with greater translational efficiency to the advantage of infecting viruses. This is supported both by our previous observation that disruption of the CCR4-NOT complex leads to longer poly(A) tails on human mRNAs and hinders HCMV infection (25) and the separate demonstration using ribosome profiling of superior translational efficiency of HCMV mRNAs over host mRNAs (58) with a mechanism of selectivity that was hitherto unexplained.

Our observations that increased poly(A) tail lengths on viral RNAs is a general feature of herpesvirus infections, regardless of subfamily, suggest that a conserved and fundamental mechanism underlies this phenomenon. One possibility is that longer poly(A) tails are preferentially installed on herpesviruses mRNAs however this would be dependent on functionally modulating PAP or PABPN1 in a manner that selectively impacts viral mRNAs. Alternatively, the hijacking of readenylation pathways could lead to preferential lengthening of poly(A) tails. This explanation would be supported by a study demonstrating that depleting the human RNA-binding protein CPEB1 led to shorter poly(A) tails on HCMV RNAs (59). We also detect the hallmark non-A incorporation by TENT4 polymerases on select herpesviral mRNAs suggesting that multiple readenylation pathways could be in operation. A third possibility is a resistance to deadenylation. Our results show that mixed tailing frequencies on host and viral mRNAs remain generally low and thus cannot explain long poly(A) tails present on most herpesviral mRNAs. Multiple herpesviruses including KSHV (60), EBV (61), MCMV (62), and HCMV (63), however, encode proteins that interact with the CCR4-NOT complex with undefined functional significance, potentially allowing herpesviruses to influence its activity or mRNA target specificity.

A simpler potential explanation for longer poly(A) tail lengths on viral RNAs could be that as a population they are simply younger than the existing pool of human transcripts in the infected cell. Arguing against this notion is our results from other virus mRNAs, including those of the nuclear DNA virus adenovirus, which do not show the same extended poly(A) tail compared to human mRNAs. Similarly, we also did not observe consistent differences in poly(A) tail length distributions between different temporal classes of HSV-1 and HCMV RNAs (with the possible exception of HSV-1 IE gene transcripts), although interpretation of these data is complicated by continued expression of RNAs from early temporal classes at late times post infection. Moreover, the median half-life of human mRNAs is estimated to a be ∼10 hours (64), thus we would expect the population of host transcripts to be largely renewed during the course of most of the infection conditions we examined.

Within the viral mRNA populations examined at the transcript level we also found transcript-specific patterns of poly(A) tail length changes over infection, suggestive of additional intersecting mechanisms. Indeed, systematic screening of viral RNA elements recently identified several from herpesvirus mRNAs that influenced reporter mRNA abundance and translation (65) and investigation of select herpesviral transcripts has identified several host and viral RNA binding proteins (RBPs; for example (66)) whose effects on poly(A) tail length are yet to be investigated. Our analysis also revealed that poly(A) tails on herpesviral ncRNAs differ dramatically in length, highlighting bespoke mechanisms. For instance, poly(A) tails on the nuclear-retained KSHV ncRNA PAN are short, in contrast to viral mRNAs and the cellular nuclear-retained ncRNA NEAT1_1. PAN RNAs have been shown to protect their poly(A) tails through interaction with U-rich sequences within its expression and nuclear retention element (ENE) (51). Notably, the modal poly(A) tail length of PAN RNAs (69 nt) is a near-perfect match for the 76nt ENE. This, and the paucity of shorter poly(A) lengths on PAN (Fig. 3b) suggests that the ENE interacts with the most 5’ part of the PAN poly(A) tail, allowing trimming from its initial ∼200 nt length, but protecting the bound region from further trimming. Among HCMV ncRNAs RNA2.7 poly(A) tails are notably longer than the others. While localised to the cytoplasm and presumed to be a substrate for deadenylases, RNA2.7 includes a CNGGN-type pentaloop that can recruit TENT4A/B enzymes to install ‘mixed tails’ that are refractory to deadenylation (26). Our analysis showed that only a small proportion of RNA2.7 molecules contain non-As, suggesting that additional mechanisms are employed to counter deadenylation. This could include recruitment of PABPC proteins to an internal A-rich sequence resulting in PABPC:PABPC interactions that could provide a mechanism to protect poly(A) tails. While only 12 adenosines are required for PABPC binding (67), a stretch of 14 adenosines interrupted by a single guanosine is found internally in RNA2.7, as well as other shorter A-rich sequences (68).

We anticipate that a consequence of retaining long poly(A) tails on viral RNAs and HCMV RNA2.7 in particular, would be to significantly sequester poly(A)-binding proteins, as well as other RBPs, and potentially to functionally deplete deadenylase activity toward cellular mRNAs. In agreement with this, AU-rich mRNAs, which are typically destabilized by mechanisms requiring CCR4-NOT, were found to be stabilized in HCMV infection in an RNA2.7-dependent manner (69). Conversely, a new pre-print suggests that sequestration of poly(A)-binding proteins by coronavirus RNA poly(A) tails accelerates the deadenylation and decay of short-tailed host mRNAs, accounting for an apparent longer tail length on the remaining population (70). Our findings in SARS-CoV-2 infected cells are consistent with these results. Which outcome prevails may be influenced by the relative abundance of viral RNAs, their tail lengths and composition, as well as virus effects on RNA decay and RBP abundance.

An open question is whether herpesviral RNAs produced during latency also retain longer poly(A) tails. While viral gene expression levels during latency varies between herpesviruses, the overall output is generally low, and for many models, only a small portion of cells are infected. However, a recent study of EBV latently infected B-cells reported mean poly(A) tail length values on viral mRNA to range from 81 - 125 nt, and on probable viral ncRNAs from 105 - 300 nt (40), indicating that long poly(A) tails on herpesviral RNAs is a consistent feature for which intrinsic sequence features may be sufficient. Applying DRS to other herpesviral latency model remains technically challenging and will likely require the development of tailored enrichment protocols.

Finally, while a number of unresolved questions remain, it is clear that nanopore DRS methodologies offer a simple and expedient way to analyse poly(A) tail lengths across individual RNAs without the biases associated with short-read sequencing methodologies. Successfully combining DRS with cellular fractionation and metabolic labelling approaches would enhance this approach further, enabling users to differentiate between effects on initial poly(A) tail synthesis and its subsequent metabolism.

In summary, we have shown that long poly(A) tails are a feature of herpesviral RNAs but that this does not broadly extend to other virus families. We further validated previous observations that increased rates of mixed-tailing is a feature of HCMV infections but argue the low frequency at which this occurs (and the variation among herpesviruses) demonstrates this is not sufficient to explain how HCMV or herpesvirus RNAs in general, broadly resist deadenylation and that additional critical mechanisms for modulating poly(A) tail lengths on viral RNAs remain unidentified.

## Acknowledgements

DPD is supported by a German Centre for Infection Research (DZIF) Associate Professorship and also receives funding from the Deutsche Forschungsgemeinschaft (DFG, German Research Foundation) under Germany’s Excellence Strategy - EXC 2155 - project number 390874280 (https://www.resist-cluster.de/en/). HB is supported by the Medical Research Council (MR/Z505523/1) and the Academy of Medical Sciences (SBF008\1027). EL, PC, and CJ were supported by the Hannover Biomedical Research School (HBRS) and the Center for Infection Biology (ZIB). CJ was funded by the Deutsche Forschungsgemeinschaft (DFG, German Research Foundation) project number 405772731 (GZ: VI 761/1-1). KK was funded by the Deutsche Forschungsgemeinschaft (DFG, German Research Foundation)—SFB 900/3—project number 158989968 to AV-B (TPB9) (https://www.mh-hannover.de/sfb900.html). ACW is supported by NIAID grants R01AI176335 and R01-AI170583. REW, WW, and MF are supported by the Medical Research Council (MR/Z505444/1). We thank Theo M. Bestebroer (Erasmus MC) for providing MPXV strain NL001-2022. The iSLK.219 cells were kindly provided by Eleonora Naimo and Thomas F. Schulz (Hannover Medical School).

---

**Figure S1:** Further analyses of mixed tailing in human and herpesviral mRNA poly(A) tails. **(a)** Breakdown of nucleotides present in mixed tails for a synthetic RN7SK RNA with a clean tail and two additional biological replicates for HCMV-infected cells harvested at 72 hours post infection (siCTRL), (a) cellular mRNA and (b) HCMV mRNAs. **(b)** Breakdown of nucleotides present in mixed tails of two additional biological replicates HCMV-infected cells harvested at 72 hours post infection. **(c)** Position specific frequency of different non-A nucleotides in the poly(A) tail of selected transcripts and timepoints. Poly(A) tails have been aligned by their 3’ end, positions are defined by their distance of the ultimate 3’ nucleotide. The three 72h datasets have been pooled. Plotted frequencies are calculated by the number of specific non-adenosine residues with a certain distance d to the 3’ end relative to the total number of reads with a poly(A) tail length ≥d, averaged with a sliding window of 5 nucleotides.

**Figure S2:** Transcript- and nucleotide level incorporation of non-adenosine nucleotides in human mRNA poly(A) tails predicted by *ninetails*. Frequency of non-adenosine residues per 1000 nucleotides in human mRNAs across different timepoints. A per mRNAs coverage filter of ≥30 was applied, only transcripts that fulfill this criterion across all samples are included. Boxplots whiskers depict 1.5 * IQR, the mean is shown by the green dot. Kruskal-Wallis rank sum test was applied on all groups for every of the three nucleotides, followed by Dunn’s test with Bonferroni correction. Significant adjusted p-values (≤0.05) are marked above the boxplots, complete Z-scores and adjusted p-values can be found in Table S5.

