## Supplementary figures and images for "Extended poly(A) tails are a shared feature of herpesvirus mRNAs"

### Supplementary Figure 1

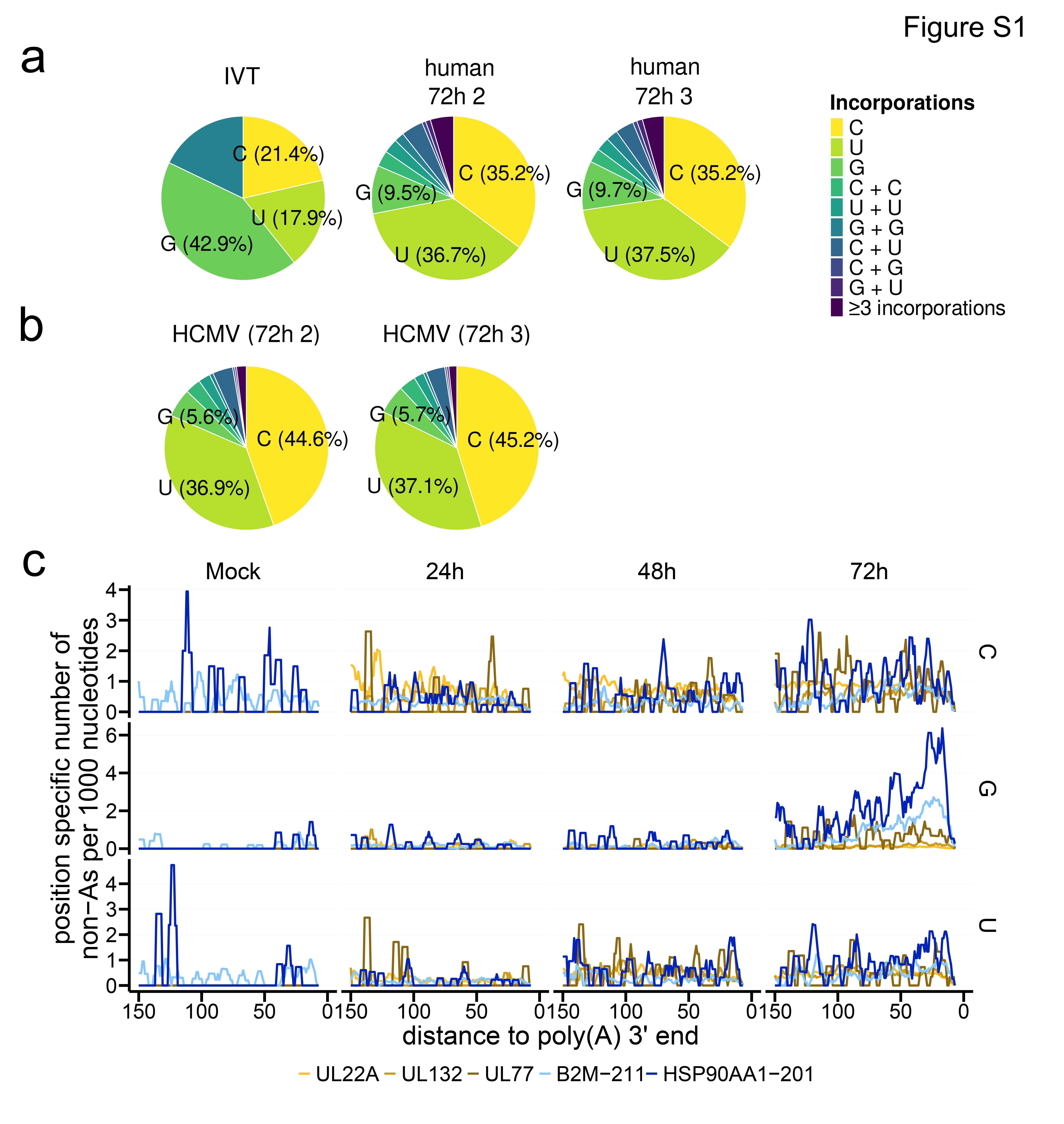

### Supplementary Figure 2

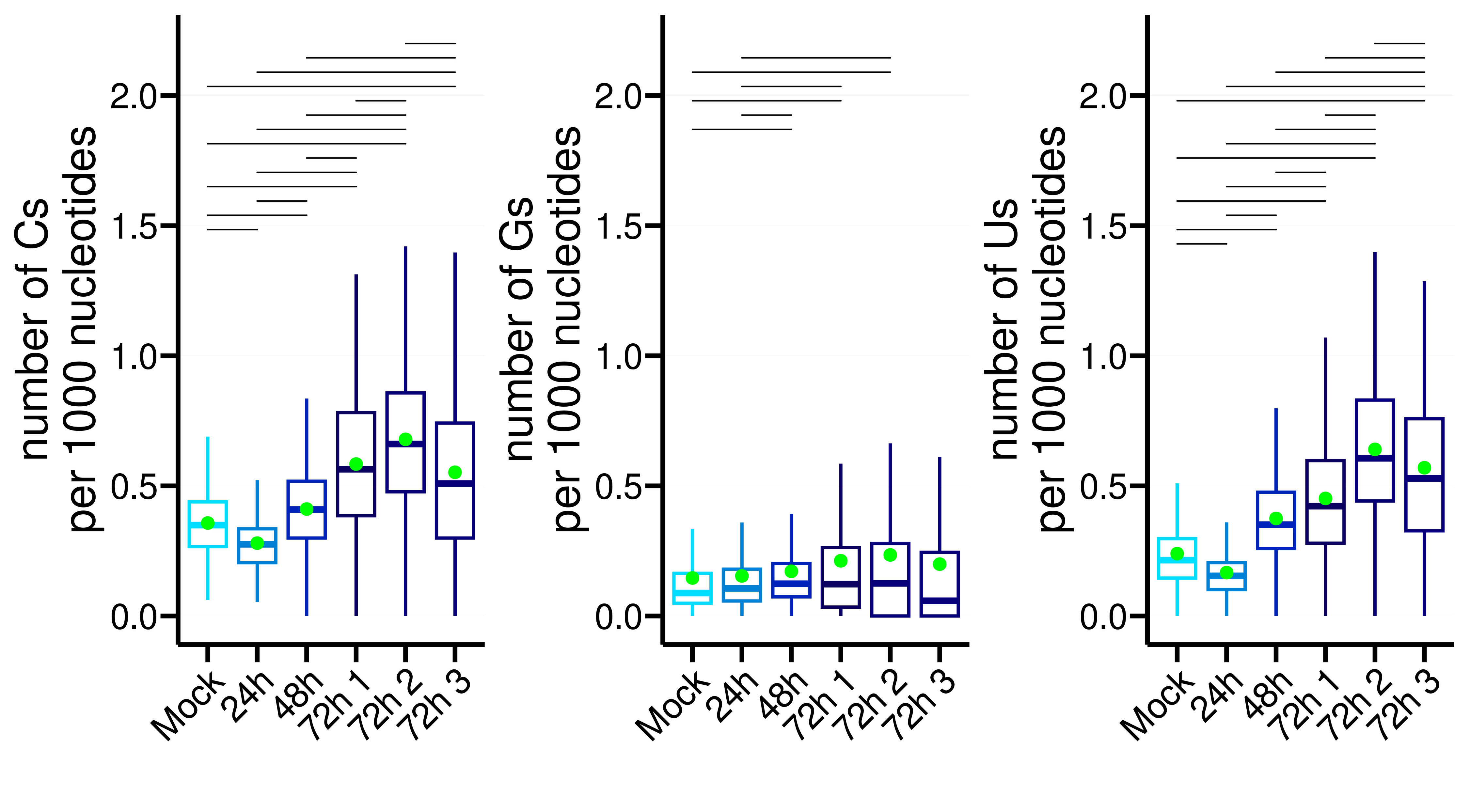
